# Self-sustained Planar Intercalations due to Mechanosignaling Feedbacks Lead to Robust Axis Extension during Morphogenesis

**DOI:** 10.1101/837450

**Authors:** Samira Anbari, Javier Buceta

## Abstract

Tissue elongation is a necessary process in metazoans to implement their body plans that is not fully understood. Here we propose a mechanism based on the interplay between cellular mechanics and primordia patterning that results in self-sustained planar cell intercalations. Thus, we show that a location-dependent modulation of cell mechanics due to positional information leads to robust axis extension. To illustrate the plausibility of this model, we use different experimentally reported patterning mechanisms in simulations that implement mechano-signaling feedback. Our results suggest that robust elongation relies on a trade-off between cellular and tissue strains that is orchestrated via the cleavage orientation. In the particular context of axis extension in Turing-patterned primordia we report that the combination of different directional cell activities lead to synergetic effects. Altogether, our findings help to understand how the emerging phenomenon of tissue elongation emerges from the individual cell dynamics.

## INTRODUCTION

During development, the initial spherical symmetry of the zygote undergoes complex changes in size and shape to form different tissues/organs and implement the body plan [1]. In that context, axis elongation, or more generically anisotropic growth, is a key morphogenetic geometric transformation that relies on the regulation of cellular activities due to the interplay between signaling and cell mechanical properties [2–8]. During axis elongation, signaling events establish planar polarity patterns at the tissue level that feed back into the cellular dynamics, e.g. oriented mechanical responses. A notable example is the convergent-extension (CE) phenomenon due to cell intercalation events [9–12]. In other cases, a directed developmental expansion is achieved by translating polarity into differential growth events, oriented divisions, and/or active migration [13–15]. More recently, this problem has been addressed from the viewpoint of the changing physical properties of tissues [16]. Recent relevant examples include evidence that shows that during the vertebrate body axis extension tissues undergo a jamming transition from a fluid-like behavior to a solid-like behavior [17]. Yet, open questions remain. On the one hand, it is not clear how instructive signals arising from primordia patterning are effectively converted into a non-equilibrium cellular dynamics that robustly sustains tissue extension and modulates their material-like properties spatiotemporally. On the other hand, further research is needed to understand how different mechanisms may contribute, cooperatively, to achieve robust anisotropic growth.

Some of these questions are beautifully illustrated during the limb bud outgrowth, a model system in morphogenesis to understand patterning and the directed developmental expansion of tissues [18, 19]. In that regard, some models have considered the proliferation gradient hypothesis [20–23]. These models are supported by the demonstrated existence of a fibroblast growth factor (FGF) gradient that has its source at the apical ectodermal ridge (AER) [1]. However, while there is experimental evidence of a spatial modulation of the cell proliferation rates, it has been shown that this mechanism is not enough to generate a significant distal limb bud outgrowth [24]. Thus, it has been suggested that limb elongation must be driven by “other”, or additional, directional cell activities. Following these ideas, recent models have proposed that limb outgrowth relies on a CE mechanism based on the existence of an anisotropic filopodial-tension [25]. Notably, this model is able to resolve a conundrum that is repeatedly found during axis elongation by CE (and observed during the limb bud outgrowth): cells elongate perpendicularly to the direction of the axis extension [24]. Still, how the pattern of gene expression observed in the limb bud modulates the cellular mechanics to generate such behavior in a sustained way is not understood.

Here we propose a framework to understand tissue elongation during morphogenesis, in particular that of the limb bud. Our model relies on the interplay between cellular mechanical properties and the patterning due to signaling that provides positional information to cells within a primordium. Here we show that such feedback, when combined with cellular growth and division, leads to auto-catalytic cell intercalations that can sustain tissue elongation robustly. We illustrate our proposal by means of numerical simulations of growing tissues using a vertex model [26, 27] that allows for mechano-signaling feedback. We show the applicability of our proposal to different situations by simulating two distinct developmental patterning mechanisms: the French Flag model [28] and the Turing instability [29].

## RESULTS

### From Cell Signaling to Tissue Elongation: an Auto-Catalytic Cell Intercalation Mechanism

Here we propose a mechanism that explains how an equilibrium, stationary, pattern of planar polarity at the tissue level, as determined by the cell signaling activity, is translated into a non-equilibrium process able to elongate tissues. Our proposal is sketched in Fig. 1. Gene regulation and long/short-range cell-cell communication lead to a planar polarity pattern (Fig. 1A). Downstream signals further refine the pattern and provide positional information to cells in terms of distinct domains that determine cellular identities (Fig. 1B). If cell identity confers distinct mechanical properties that promote intermingling among cells from different domains, then cells by domain boundaries intercalate to minimize their energy (Fig. 1C). As for the cellular division process, cell growth and intercalation-induced stretching coupled to the Hertwig rule (cleavage orientation perpendicular to the longest cell axis) set the preferential orientation of cleavage planes parallel to domain boundaries [30, 31]. Such bias in terms of the elongation and division orientation has been experimentally reported in a number of developmental processes including limb development [24, 32–34]. Following division, identities, and hence mechanical properties, of daughter cells are reassigned depending on their position within the tissue. Dynamical assignment of cellular identities depending on their locations in a morphogenetic field is common during development, e.g. [35]. In that regard, experimental evidence about the dynamic establishment of cellular identities in the case of the limb bud primordium comes from micromass cultures where it has been shown that up-regulation, or down-regulation, of the skeletal marker *Sox9* depends on the relative positions of cells within the tissue [36]. This feedback between intercalation, division, and dynamic identity switching results in an auto-catalytic cell intercalation mechanism at the domain boundaries that leads to a self-sustained CE process (Fig. 1C). Self-sustained intercalations cause tissue extension while cells elongate perpendicularly to the extension axis as schematically represented by the polar histogram cartoons in Fig. 1D. See Methods for details about the mathematical formalization and the computational implementation of this mechanism. We stress that cell intercalations that rely just on the differential adhesion hypothesis (DAH) cannot generate tissue elongation. It that case, distinct mechanical properties of cells (canonically promoting phase separation of homotypic populations [37–39]) are *inherited*. As a result, a transient CE may occur depending on the initial configuration but, in the long term as cells grow and divide, it leads to isotropic tissue growth (Fig. S1 and Movie S1). Here, instead, we propose a modulation of the mechanical properties such that cellular *affinities* (i.e., energetic costs) are assigned, dynamically, by positional information (rather than inherited).

**Figure 1.**
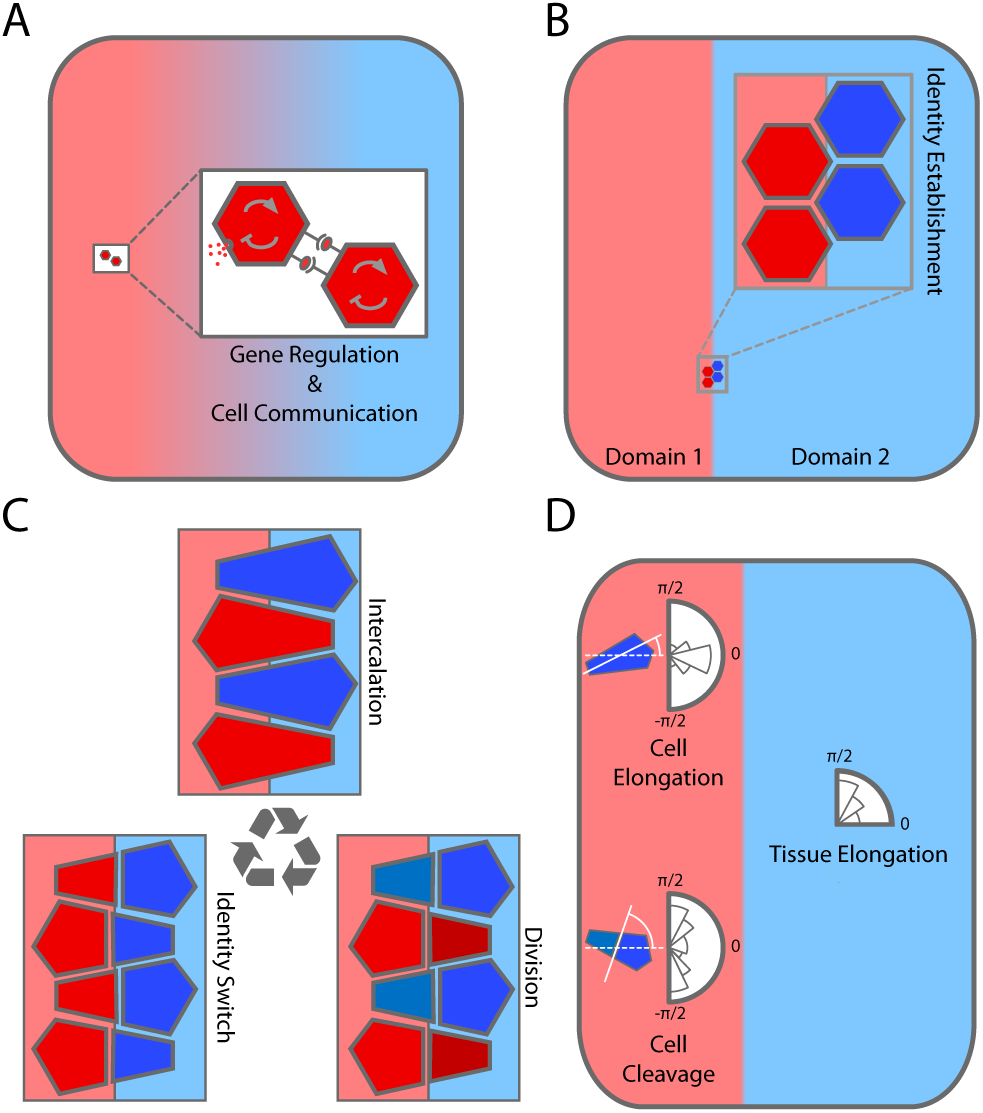
A mechanism to translate tissue planar polarity into self-sustained CE. Signaling and communication pattern the tissue (**A**) and establish positional information domains that provide cellular identity (**B**). **C:** If cell identity implies distinct mechanical properties that promote cell intermingling, it leads to an auto-catalytic intercalation mechanism (see text). **D:** Orientation features of cellular processes, such as elongation or cleavage, indicate that cells elongate along an axis perpendicular to the domain boundary, but the resulting CE arising for intercalations extends the tissue along a direction parallel to the domain boundary.

### Auto-Catalytic Cell Intercalation Induces Axis Elongation in Morphogen Patterned Tissues

As a proof of concept about the functionality of the proposed auto-catalytic cell intercalation mechanism, we simulated a tissue patterned by a morphogen gradient (Methods) [40]. Our results showed that regardless of the growth and division activity of cells, a stationary gradient profile is established (Fig. S2 and Movie S2). Positional information was provided to cells following the French Flag model paradigm that sets domains of dynamic cellular identities depending on morphogen concentration thresholds (Methods) [28]. We assumed distinct mechanical properties of cells in terms of adhesion (i.e., line tension in our vertex model implementation) such that there is an increased affinity between cells from different domains that promotes cell intermingling by domain boundaries (Methods). Figure 2A (Movie S3) shows that a noticeable elongation of the tissue is achieved in contrast to control simulations where cell adhesion is not modulated by the morphogen signal (Movie S4). For a precise mathematical definition of the elongation ratio see Methods. In these *in silico* experiments, we implemented the Hertwig rule for cell divisions; yet including variability as experimentally reported [41] (Methods).

**Figure 2.**
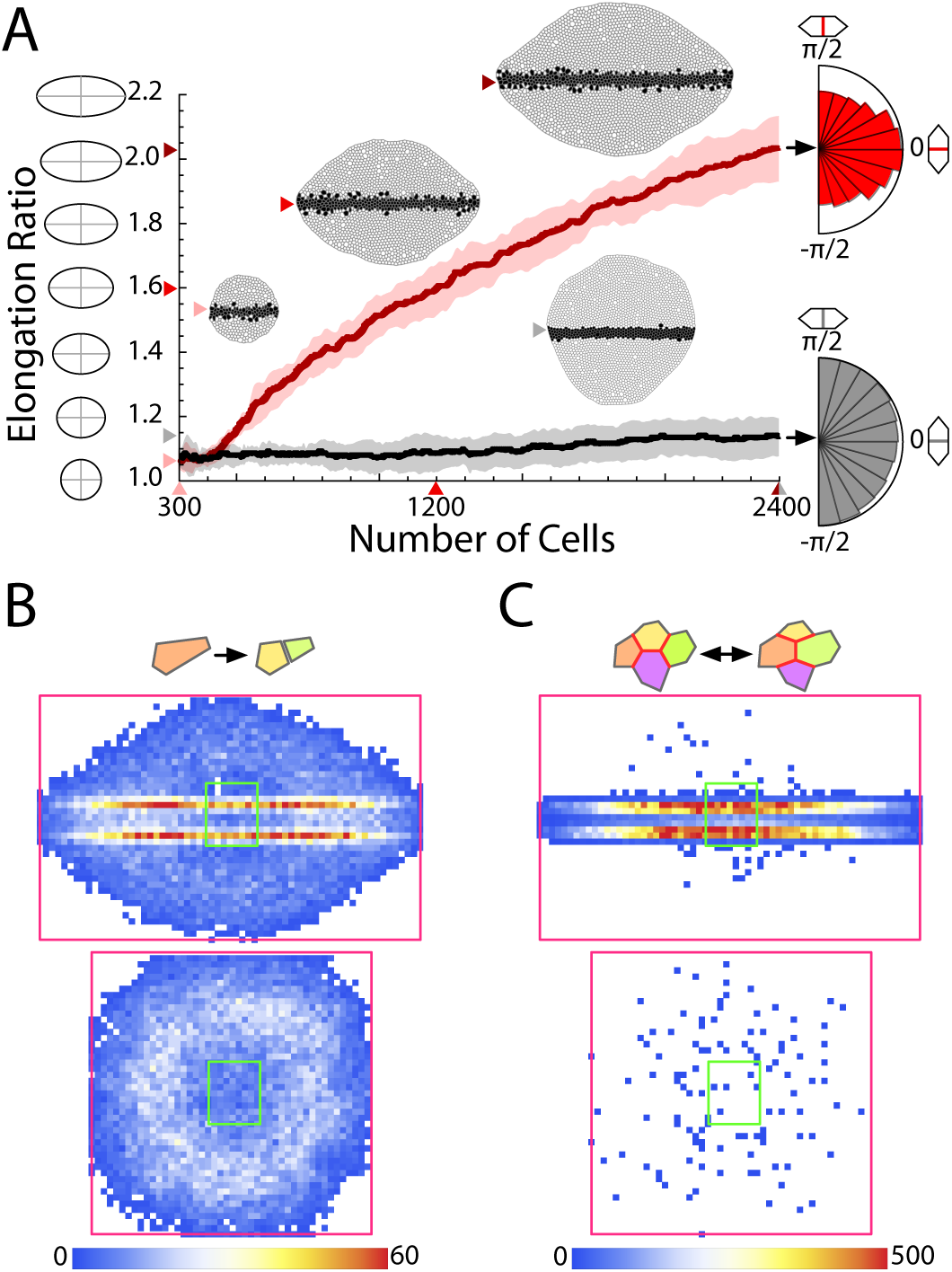
Coupling between tissue patterning and cell mechanics leads to robust elongation. **A:** Elongation ratio as a function the number of cells in tissues patterned by a morphogen gradient (*in silico* experiments). Red stands for the case when cell mechanics and patterning are coupled. Black refers to control simulations (adhesion not modulated by the pattern). Results from ten simulations: solid lines indicate the average and the shading the standard deviation band. Values of average elongation, number of cells, and snapshots of representative simulations as indicated by the colored arrows. The cumulative polar histograms of cleavage events (right) reveal that cells preferentially elongate perpendicular to the extension axis when the auto-catalytic intercalation mechanism applies. **B:** Cumulative density histograms of divisions (ten simulations). The green/magenta squares indicate the initial/final bounding boxes that delimit the tissue size. Intercalation-induced cell stretching (top) promotes cell divisions at the domain boundaries. **C:** Cumulative density histograms of T1 transitions (ten simulations). Auto-catalytic intercalation (top) provides fluidity to the tissue as evidenced by its active remodeling at domain boundaries.

Hence, the quantification of the cleavage orientation is as a proxy for the cellular elongation direction. In that regard, our results indicate that cells preferentially elongated perpendicular to the tissue extension axis due to the intercalation events (Fig. 2A). If cell adhesion is not modulated by the morphogen signal we observed a randomized cellular elongation as expected (Fig. 2A). Intercalation induces cell stretching that in turn promotes cell division as experimentally reported [42]. To test this possibility, we quantified the cumulative density of division events and found that cells indeed divided more actively at domain boundaries where intercalation is more operative (Fig. 2B). That is, the material-like additive properties of the tissue are enhanced by the auto-catalytic mechanism. In contrast, when adhesion is not modulated by the tissue patterning, cell division locations are not correlated with the position of the domain boundaries. We also observed that inner cells eventually divided less often due to the pressure increase [43]. Finally, to evaluate the plasticity (fluidity) of the tissue, we computed the accumulated density of T1 topological transitions (Fig. 2C). In that regard, we found that auto-catalytic intercalations largely increased the cellular activity, thus making the tissue more fluid.

### Robust Elongation Relies on a Trade-off between Cellular and Tissue Stresses

In order to clarify more precisely the role played by oriented cell divisions during axis extension, we performed additional *in silico* experiments where we tested distinct alternatives to the Hertwig rule: random orientation of the cleavage plane and divisions following the opposite of the Hertwig rule (cleavage plane parallel to the longest cell axis). As in the case of the Hertwig rule, we included variability (i.e., noise) in the cleavage orientation. We found that if the cleavage orientation followed the opposite of the Hertwig rule then, while the auto-catalytic intercalation mechanism still applies, the magnitude of the axis extension is lessened and the overall shape of the tissue is very irregular, Fig. 3A-B. Random cleavage orientation implied an intermediate situation where elongation is achieved, but the tissue shape developed some irregularities, Fig. 3A-B. Interestingly, in the case of the “opposite” rule, one would expect a cleavage statistics that would be the opposite to that found in Hertwig; yet we found that division planes parallel to the extension axis are still predominant (see polar histograms). To investigate this phenomenon, we examined the cellular dynamics within the different positional information domains of the tissue. Our results indicate that auto-catalytic intercalation generates cellular stresses in the domain where this mechanism is more active (central domain) that contributes to elongate the cells perpendicularly to the extension axis, Fig. 3C.

**Figure 3.**
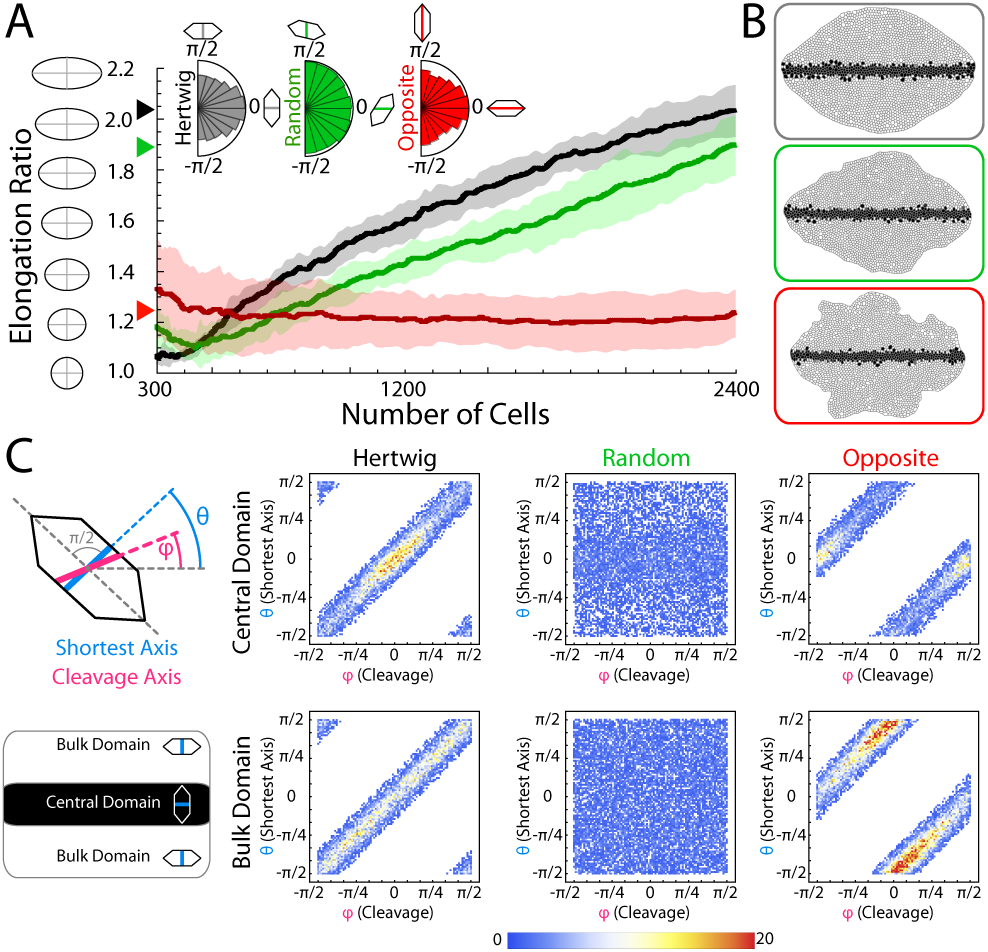
Effect of cleavage orientation. **A:** Tissue elongation as a function of the number of cells for different cleavage dynamics (ten simulations): solid lines stand for the average and the shadings for the standard deviation bands. The polar histograms of cleavage orientations (inset) are a readout of the cellular geometry when the Hertwig rule (black) or its opposite (red) apply but not in the case of random cleavage (green). The Hertwig rule leads to a systematic elongation. **B:** Final snapshots of representative simulations depending on the cleavage dynamics (color codes as in **A**). If cells do not divide perpendicularly to their longest axis, then the tissue develops irregularly. **C:** Analysis of the correlation between cellular geometry prior to division (as represented here by the shortest cell axis, *θ*) and the cleavage orientation (*φ*) in different tissue domains by means of cumulative density histograms.

As a result of the cellular intercalation the tissue elongates, that in turn generates stresses in the cells of the “bulk” domain that promotes their elongation *along* the extension axis, Fig. 3C. This orthogonal orientation of cellular geometries in different domains is, in fact, more clearly revealed when the “opposite” mechanism applies. Altogether, these results suggest that the interplay between cellular and tissue stresses, when coupled through oriented cell divisions following the Hertwig rule (even in the presence of noise), is instrumental in generating a robust axis elongation.

### A. Axis Extension in Turing Patterned Tissues Depends on Synergetic Mechanisms

As shown above, the mechanism introduced here implies an instructive role of signaling cues to determine the elongation axis: tissues extend along the direction of the boundary domains determined by planar polarity. In that regard, in primordia patterned following the French Flag model the elongation axis is set by the signaling center from which the morphogen is produced and diffuses out. This raises the question of whether the proposed mechanism applies to more complex patterning situations that need auxiliary mechanisms to establish planar polarity at the tissue level. To that end, we studied Turing patterned tissues. Developmental examples of the latter include animal coating [44], tooth primordium patterning [45], the rugae spacing in the mammalian palate [46], and a case that is particularly relevant in the context of tissue elongation: limb bud outgrowth [36, 47].

Turing instabilities set distinct domains of expression in tissues. However, the patterns always display some level of rotational symmetry. Different ideas have been suggested to achieve stripe alignment (i.e., a symmetry-breaking event) in the context of Turing patterns [48]. While it has been argued that all the proposed mechanisms produce robust stripe alignment, we found in the *in silico* experiments that, when tissues are subjected to cellular growth/division, the diffusivity-modulation mechanism due to the activity of a morphogen released from a cellular population consistently leads to pattern alignment (Fig. 4, Methods, Discussion). Yet, we stress that the applicability of the auto-catalytic intercalation phenomenon is independent of the auxiliary mechanism that promotes stripe alignment. In the particular context of the limb bud, FGF, released from the AER, would play the role of the morphogen setting the planar polarity pattern that induces pattern alignment [36]. In addition, there is experimental evidence showing that FGF stimulates outgrowth and cellular proliferation [49]. Thus, we tested how axis extension in Turing patterned tissues depends on synergistic interactions between different mechanisms.

**Figure 4.**
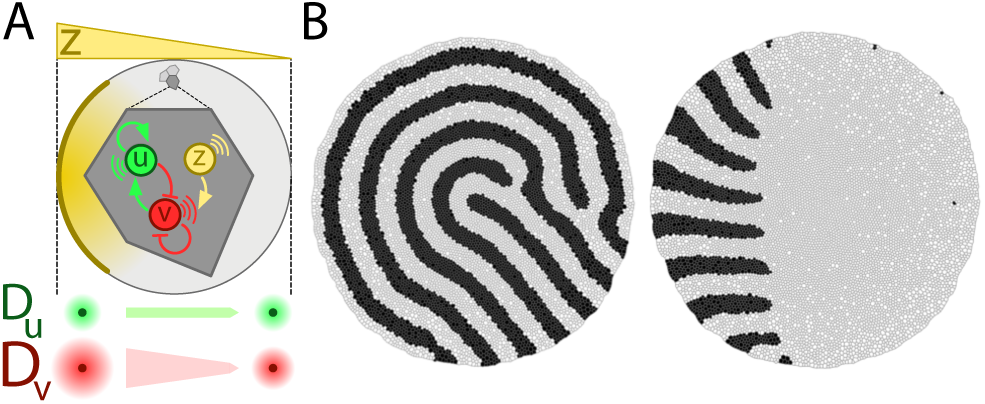
Stripe alignment mechanism in growing tissues. **A:** In tissues where cells actively grow/divide, if each cell is driven by a regulatory network that just involves an activator, *u*, and an inhibitor, *v*, then the resulting Turing pattern displays rotational symmetry (panel **B** left). If an additional species, *z*, is released from the “tip” (left side of the tissue in this example) and set a polarity gradient such that the diffusivity of *v* is spatially modulated, then stripes align following the gradient directionality (panel **B** right). **B:** Final snapshot of simulations without (left) and with (right) diffusivity modulation (constant cellular adhesion). The black (white) cellular domains account for regions where *u* > *v* (*v* > *u*). Since diffusive transport relies on tissue topology (Methods) we avoided a possible bias in patterning by using in both simulations the same random sequences that determine the variability of cellular growth/division in order to reproduce the same cellular growth/division events and cell/tissue topologies.

Figure 5A (Movie S5) shows that a combination of a spatial modulation of the cellular proliferation rates (cell cycle speed proportional to the morphogen signal released from the tip, Methods) and the auto-catalytic intercalation mechanism leads to a robust (less variability), and fast, axis extension. Motivated by prior studies that showed that a modulation of proliferation rates is not enough to generate a significant distal limb bud outgrowth [24], we implemented in our simulations a very “mild” modulation: ∼ 2% increment of the elongation ratio with respect to control simulations that lack auto-catalytic intercalations and a modulation of proliferation rates (Fig. 5A and Movies S6-S7, see also control Movie S8). Yet, when combined with the auto-catalytic intercalation mechanism, the elongation was boosted by ∼ 14%. Also, as revealed by Fig. 5B, cells elongated perpendicularity of the tissue expansion direction as long as the auto-catalytic intercalation mechanism applies.

**Figure 5.**
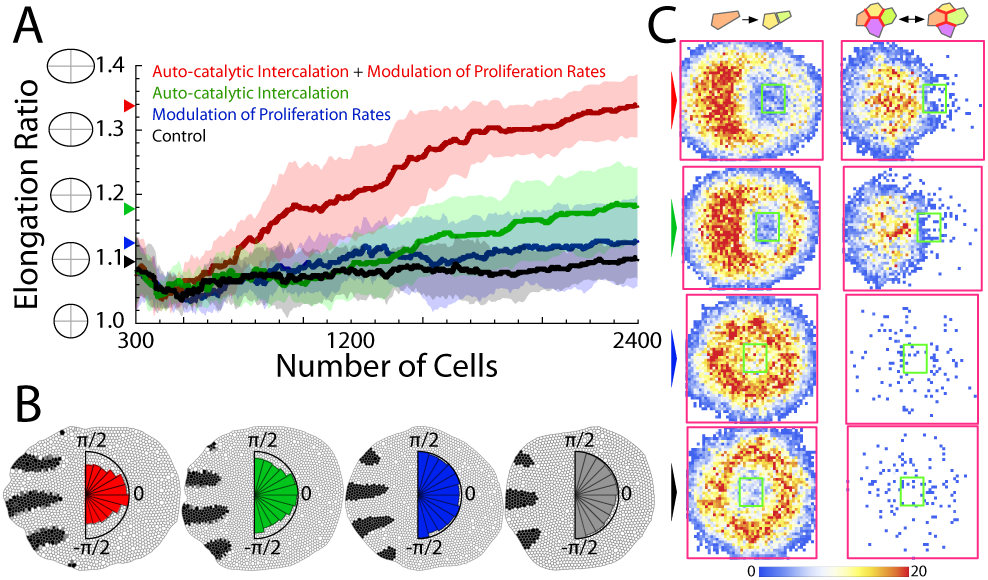
Elongation in Turing patterned tissues. **A:** Comparison of tissue elongation as a function of the number of cells using different mechanisms (ten simulations): solid lines stand for the average and the shading for the standard deviation band. In all cases, cell cleavage follows the Hertwig rule. In the control simulation, the mechanical properties of cells are independent of patterning (Supplementary Methods). The final average tissue elongations are as indicated by the arrows. **B:** Representative snapshots of simulations (same number of cells) and cumulative polar histograms of cleavage orientations (color codes and in **A**). The black (white) cellular domains account for regions where *u* > *v* (*v* > *u*). When auto-catalytic intercalation applies cells elongate perpendicularly to the direction of axis extension. **C:** Cumulative density histograms of cell division events (left) and T1 transitions (right) depending on the elongation mechanism (color codes as in **A**). The green/magenta squares indicate the initial/final bounding box delimiting the tissue size.

As in the case of the French Flag model, we observed that division events are promoted in the intercalation region (Fig. 5C). Yet, we found a less structured (i.e., digitate-less) distribution in agreement with limb bud outgrowth data [24]. As for the analysis of the topological remodeling of the tissue (T1 transitions), as a proxy for its fluidity, it revealed a clear proximal-distal (right-left in the figure) gradient when the auto-catalytic intercalation mechanism applies such that there is more plasticity in the growing tip. We also observed a less structured T1 pattern in comparison with the French Flag model simulations. Finally, as for the effect of the division dynamics we found, in agreement with the French-Flag model simulations, that either random or “opposite” cleavage dynamics leads to growth irregularities in the tissue and large variability, i.e., lack of robustness (Fig. S3).

## DISCUSSION

Here we propose a framework to understand how the interplay between patterning and mechanics leads to axis extension. Our approach provides a simple, plausible, mechanism to understand how the tissue-level planar polarity patterns that originate from cell signaling and communication set an original symmetry breaking that feeds back to the cellular mechanics to produce sustained anisotropic growth elongation via auto-catalytic intercalations. This mechanism is based on some basic assumptions that have been experimentally observed in morphogenetic processes. First, cell identities can be dynamically assigned depending on the locations of cells within a primordium following the positional information paradigm. Second, distinct identities imply cell affinities (adhesion) that promote intermingling. Third, cell cleavage follows the Hertwig rule such that cells divide perpendicular to their longest axes. Importantly, our results show that these premises lead to a non-equilibrium cellular dynamics that, without any further assumptions, is able to explain the reported directional activities of cells during CE: intercalating cells predominantly elongate perpendicular to the axis extension direction and their divisions are oriented parallel to that axis. Moreover, we have shown the existence of a trade-off between cellular and tissues strains that contribute to a robust extension dynamics.

To illustrate the applicability of this mechanism we have used two patterning cases that are found during development: tissues patterned by either morphogen gradients or by a Turing instability mechanism. Our simulations are performed using a vertex model approach that allows feedback between mechanical and signaling cues and, to check the robustness of our proposal, we have included different sources of variability such as noise in the cleavage orientation and in the cell cycle length. Our proposal does not aim at explaining in detail the elongation of a specific primordium but rather to show a generic mechanism. Still, we believe that our results are particularly applicable, and relevant, to the case of limb bud outgrowth since our findings are in agreement with the behavior found experimentally in qualitative terms. Moreover, we have shown, to the best of our knowledge for the first time, how the digitate pattern develops using *in silico* experiments with a realistic cellular dynamics. An important implication of our results in the context of the limb bud is to reconcile data in terms of the possible mechanisms underlying its outgrowth. Thus, we have shown that the synergistic interaction between auto-catalytic intercalations and spatially modulated proliferation rates leads to robust elongation. Our data suggest that the former is the main driver of elongation and the latter, while not being able to explain tissue extension, boosts its effect.

Our proposal could also provide insight into the recently reported fluidization during vertebrate body axis elongation [17]. In that context, it has been shown that there is a larger tissue remodeling at the extending mesodermal progenitor zone and yet, the analysis of the orientation neighbor exchanges revealed that no systematic alignment contributes to the elongation of the body axis. In that regard, here we have shown how patterning can promote gradients of tissue remodeling during elongation and, in fact, the directionality of neighbor exchanges is, counterintuitively, opposite to the extension direction. In that sense, the framework that we present here could help to understand how primordia patterning conditions the asymmetry of tissue remodeling activity.

As a matter of discussion, here we have assumed that all the mechanical and biological interactions of the cells are described adequately by a 2D model in a planar geometry. This over-simplification is standard in the field and can possibly provide a plausible, yet basic, understanding of tissue remodeling. However, recent discoveries about the cellular behavior in 3D environments when tissues are subjected to some level of curvature, point towards an intriguing and important role of *spatial* T1 transitions [50]. In that regard, to what extent the framework presented here depends on the 3D structure of the cells is an interesting subject for further studies.

In conclusion, we have presented a model based on hypotheses that seemingly connects the ideas of primordia patterning due to gene activity with oriented cellular activities to lead to asymmetric tissue growth. Therefore, our study paves the way to better understand shape regulation during morphogenesis.

## METHODS AND MATERIALS

### Tissue simulations

Our approach is based on the vertex model originally developed by Nagai et al. [51], and further adopted to model epithelial tissues by other authors, e.g., [52, 53]. The model takes into account three energetic contributions for each cell vertex *i*:

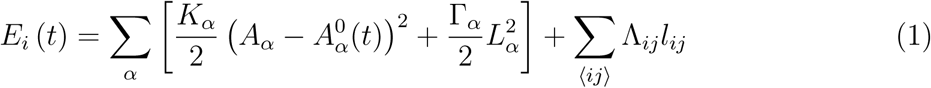

index *α* corresponds to a cell, while *i* and *j* represent adjacent vertices sharing a connecting edge. The first term (r.h.s.) stands for the elastic energy of cells caused by the difference between the *actual* cell area *A*_*α*_ and the *preferred* cell area 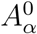 (the area that the cell would like to have due to the cytoskeleton structure in the absence of the stresses associated with the adhesion and cortical tension). The second term, proportional to the squared cell perimeter, *L*_*α*_, describes the mechanical tension related to the elastic contraction of an actomyosin cortical ring. Finally, the third term describes the adhesion energy: Λ_*ij*_ being a line tension coefficient that weights the interaction between two cells (that can be either positive or negative), and where *l*_*ij*_ represents the length of the edge connecting neighboring vertices, *i* and *j*. Based on this model, the cell packing geometries are determined by minimizing the total energy of the system which leads to a mechanical force balance where **F**_*i*_ = −∇*E*_*i*_. Under the assumption that inertia is negligible, the dynamics of cell vertices satisfies the equation of motion, 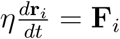 (*η* being a drag/viscous coefficient). See [53, 54] for additional details.

In our simulations we used the following dimensionless parameter values *K* = 1 and Γ = 0.02 (all simulations), Λ = ±0.05 (simulations of DAH mechanism, Fig. S1), Λ = 0.05 (simulations of the morphogen gradient profile, Fig. S2, and of Turing stripe alignment formation, Fig. 4B), for all other simulations, Λ ranges between 0 to 0.1 depending on the tissue domain see implementation of the signalling-mechanics feedback below (Auto-catalytic cell intercalation). We imposed a value for the line tension for cell edges facing the tissue exterior of Λ = 0.2. The latter promotes a circular shape of the tissue and helps to highlight that elongation or tissue deformation is due to the cellular dynamics and not to other effects.

As for the implementation of the cell cycle and the cellular growth, the cell cycle duration, *τ*, is a stochastic variable that satisfies *τ* = *ϵt*_*det*_ + (1 − *ϵ*) *t*_*sto*_ where *t*_*det*_ is a deterministic time scale that accounts for the average cell-cycle duration in the absence of mechanical stress and *t*_*sto*_ is a random variable exponentially distributed with a probability density 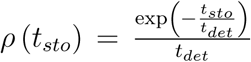. The parameter *ϵ* weights the stochasticity of the cell-cycle duration (0.8 in our simulations). In our simulations ⟨*τ*⟩ = 1.5 · 10^3^ (dimensionless). If a proliferation gradient applies due to signaling (e.g. FGF), we simulated this effect by modulating the cell cycle duration ⟨*τ* ⟩ by the morphogen concentration (see details below). Cellular growth is implemented using a piece-wise dynamics that prescribes the following growth of the (dimensionless) preferred apical cell area, 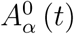: cells are quiescent up to the middle of their cell-cycle and then 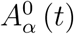 grows linearly (towards doubling) (see [53] for details). With respect to the cleavage orientation, the code evaluates the inertia tensor of cells with respect to its centre of mass assuming that a proper representation of the former is a polygonal set of rods, i.e., the cell edges. The principal inertia axes indicate the symmetry axes of the cell: the longest axis of the cell is orthogonal to the largest principal inertia axis. Cells that divide following the Hertwig rule set their cleavage plane perpendicular to the longest cell axis. In simulations where cells divide opposite to Hertwig rule or randomly, the cleavage plane is respectively parallel to the longest cell axis or random. Once the the *putative* division angle, *φ*, has been set we implement variability by using a normal distribution *N* (*φ, σ*^2^). In our simulations *σ* = 0.2 and we set bounds to the tails of the normal distribution such that the *actual* cell division lies within the interval [*φ* − 0.5, *φ* + 0.5]. Cleavage is assumed to be instantaneous in our simulations.

As for the protein dynamics, we assume each cell to be a well-stirred system where spatial effects are disregarded. Each cell may contain a number of species (proteins) with dynamics described by a deterministic differential equation (see below). Protein numbers in each cell, are calculated by a numerical integration using the Euler method and protein concentration are obtained by using the cell area at any time. Following a division event, proteins are distributed binomially between daughter cells. As for the diffusion of morphogen molecules, the diffusion operator is discretized according to the cell topology following [55].

We simulate the tissue dynamics for ∼ 5 cell cycles, yet defining two distinct temporal stages. First, starting with tissues that contain 10^2^ cells arranged in a regular hexagonal configuration, we “randomize” the topology by cell growth and cleavage events and pre-pattern the tissue by disregarding any modulation of the mechanical properties due to signaling. This stage lasts ∼ 1.5 cell cycles until the total number of cells is ∼ 300. After this transient, a second simulation stage, where modulation of mechanical properties by signaling applies, is implemented during ∼ 3.5 − 4 cell cycles until the total number of cells is ∼ 2.5 · 10^3^. All reported properties, e.g., elongation ratio, are calculated taking into account only the second simulation stage.

### French flag patterning model

We implemented the French flag patterning scheme by simulating first a signaling center from which a morphogen, *c*, diffuses out. The dynamics of the morphogen concentration for a cell *i, c*_*i*_, is prescribed following [56]:

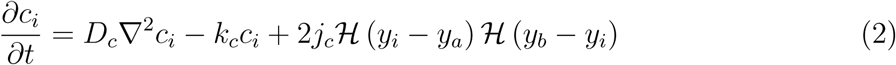

where *y*_*i*_ stands for the vertical coordinate of the geometrical center of cell *i*, ℋ (*z*) is the Heaviside step function, *D*_*c*_ is the diffusion coefficient, *k*_*c*_ the degration rate, and *j*_*c*_ the morphogen current: −*D*_*c*_*∂c*_*i*_*∂y* = *j*_*c*_ in the domain *y*_*i*_ ∈ (*y*_*a*_, *y*_*b*_). Thus, the morphogen is released from all cells with centers in the range (*y*_*a*_, *y*_*b*_). In our simulation the parameter values are (dimensionless): *D*_*c*_ = 10^−2^, *k*_*c*_ = 2.54 · 10^−3^, *j*_*c*_ = 398, and (*y*_*a*_, *y*_*b*_) = (3.5, 5). Taking into account that 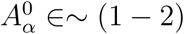, the width of the signaling center typically comprises ∼ 1 − 2 cells. Given the equation (2), if Δ*y* = *y*_*b*_ − *y*_*a*_ → 0 then the stationary concentration of the morphogen in a cell at a location *y*_*i*_ reads 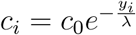 with *c*_0_ ∼ 10^5^ and 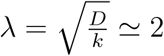 being the typical decay length of the morphogen [56]. To implement a French flag positional information mechanism, we set a morphogen threshold *c*_*t*_ = 3.5 · 10^4^ molecules/cell and defined the following rate dynamics of two putative proteins, *d*_1_ and *d*_2_ for every cell, *i*,

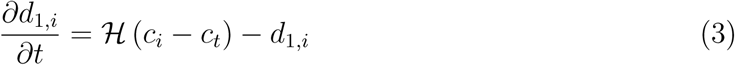

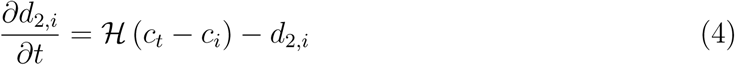

Thus, cell identities and tissue domains are characterized by a vectorial tag: central domain cells (*d*_1_, *d*_2_) = (1, 0), bulk domain cells (*d*_1_, *d*_2_) = (0, 1). Taking into account the value of *c*_*t*_ and the parameter used, the central domain has a typical width of 4 − 5 cells.

### Turing patterning model

In our simulations we used a generic reaction-diffusion model that can be mapped into an activator-substrate model proposed to describe pigmentation patterns [57] or into an activator-inhibitor model that accounts for regeneration [58]. More recently the model has been used to explored the role of the so-called protein *granular* noise due to discretization effects during patterning [59]. The model describes the concentration of two proteins, *u* and *v*, in every cell *i* that can undergo a Turing instability leading to labyrinth-like patterns with rotational symmetry:

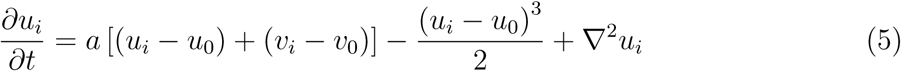

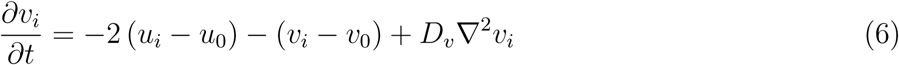

In our simulations we used the dimensionless parameters of *a* = 0.9 for all simulations and *D*_*v*_ = 9 for the simulations shown in Fig. 4B which lead to patterns around the homogeneous state *u*_0_ = *v*_0_ = 2. For details about the Turing instability condition and non-linear effects in this model see [60].

Stripe alignment was obtained by implementing the anisotropic diffusion of especies *v*. To do so, we defined a cell population with identity ℐ = *Z* at the boundary of the tissue (see Fig. 4A) that produces a morphogen, *z* (see Movies S9-S10),

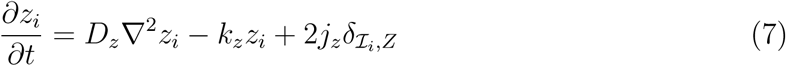

The parameters values (dimensionless) used in our simulations were: *D*_*z*_ = 0.75, *k*_*z*_ =1.25 · 10^−2^, *j*_*z*_ = 5 · 10^−2^. Under those conditions *z* takes a value of ∼ 1 at locations where ℐ = *Z*. Protein *z* modulated the diffusivity of protein *v* linearly such that 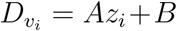with *A* = 20 (Fig. 4B, right), *A* = 13 (simulations about tissue elongation), and *B* = 4 in all cases.

Similarly to the case of morphogen patterned tissues, we defined additional putative proteins to provide a identity to cells,

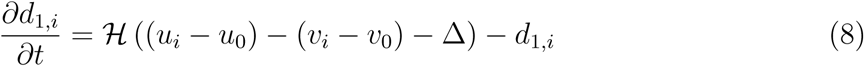

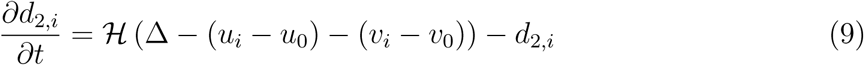

where Δ = 0.2 is a concentration threshold. That is, if (*u* − *u*_0_) − (*v* − *v*_0_) > Δ, cells are characterized by a vectorial tag (*d*_1_, *d*_2_) = (1, 0) and if (*u* − *u*_0_) − (*v* − *v*_0_) < Δ then (*d*_1_, *d*_2_) = (1, 0). Since the characteristic domain size as a function of the pattern wavelength, *l*_*c*_, is *l*_*c*_/2, and taking into account that (see [60]),

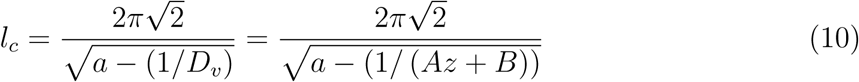

then the domains at locations where ℐ = *Z*, i.e. *z*_*i*_ ≃ 1, comprise ∼ 5 − 6 cells (Fig. 4). The patterning disappear at locations where 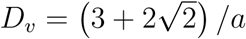 [60].

The morphogen concentration profile is further used to generate a proliferation gradient in some simulations (see text). In that case the average cell cycle duration a function of *z* is 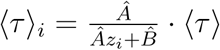 with ⟨*τ*⟩ = 1.5 · 10^3^, *Â* = 4, and 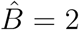. The cells cycle duration varies from 10^3^ (at locations where *z* = 1) to 3 · 10^3^ (at locations where where *z* = 0).

### Auto-catalytic cell intercalation

The patterning-mechanics interaction is implemented in our model through the putative proteins *d*_1_ and *d*_2_ that characterize dynamically the positional information depending on the underlying gene regulation that patterns the tissue. Thus, the following matrix describes the identity relation between a pair of cells neighboring *i* and *j*,

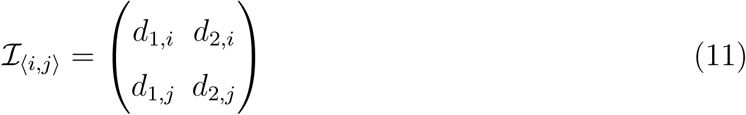

Consequently, if cells *i* and *j* belong to the same positional information domain then |det (ℐ_⟨*i,j*⟩_) |= 0 and if cells *i* and *j* belong to different positional information domain then |det (ℐ_⟨*i,j*⟩_) |= 1. In our simulations we modulated the adhesion energy between two neighboring cells by implementing the following dependence of the line tension parameter, Λ_*i,j*_, as a function of ℐ_⟨*i,j*⟩_, see equation (1): Λ_*ij*_ = Λ_0_ + *γ* |det (ℐ _⟨*i,j*⟩_)|with Λ_0_ = −*γ* = 10^−1^. As a consequence, cell intercalation is promoted at domain boundaries.

### Tissue elongation ratio

The tissue elongation ratio is computed as follows. We first estimate the center of mass of the tissue using the perimetric cell vertices. Second, we calculate the components of the inertia tensor with respect the center of mass of the tissue:

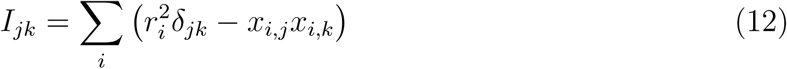

where the sum runs for all the perimetric vertices, *i*, with Cartesian coordinates (*x*_*i*_, *y*_*i*_) = (*x*_*i*,1_, *x*_*i*,2_), *r*_*i*_ is their distance to the center of the mass, and *δ*_*jk*_ is the Kronecker delta. Finally, we obtained the tissue elongation ratio by calculating the ratio of the two eigenvalues of the inertia tensor.

### Cells division and T1 transitions

The location of cell divisions is computed by collecting the coordinates of mother cell centers right before cleavage. As for T1 transitions, we registered the coordinates of the edge associated to neighbor exchanges right before, 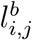, and after, 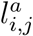, a transition. The location of a T1 transition is characterized by the intersection point of the edges 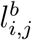 and 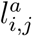.

## Supplementaty Information (SI)

**Figure 6.**
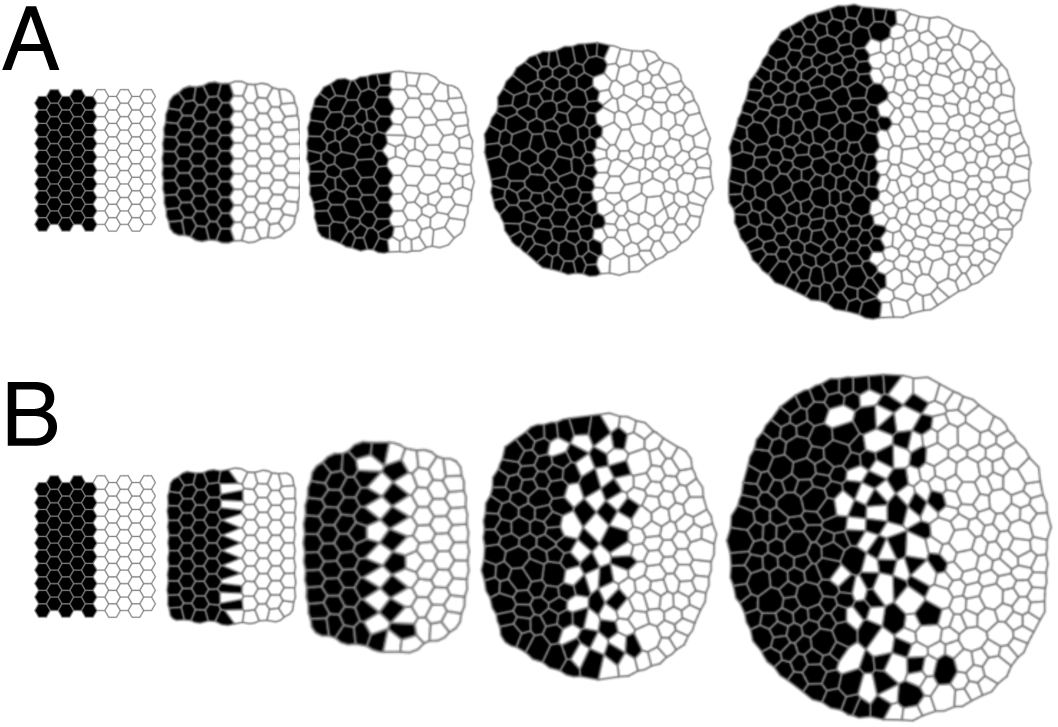
**(Figure S1)** Starting from the same initial configuration, if the cell adhesion (line tension parameter) is the same for different cell populations, **A**, then cell intercalations are not promoted. If the adhesion between cells promotes cell intermingling, **B**, cell intercalation is observed. However, the fact that the cell mechanical properties are inhereted leads to isotropic tissue growth in the long term.

**Figure 7.**
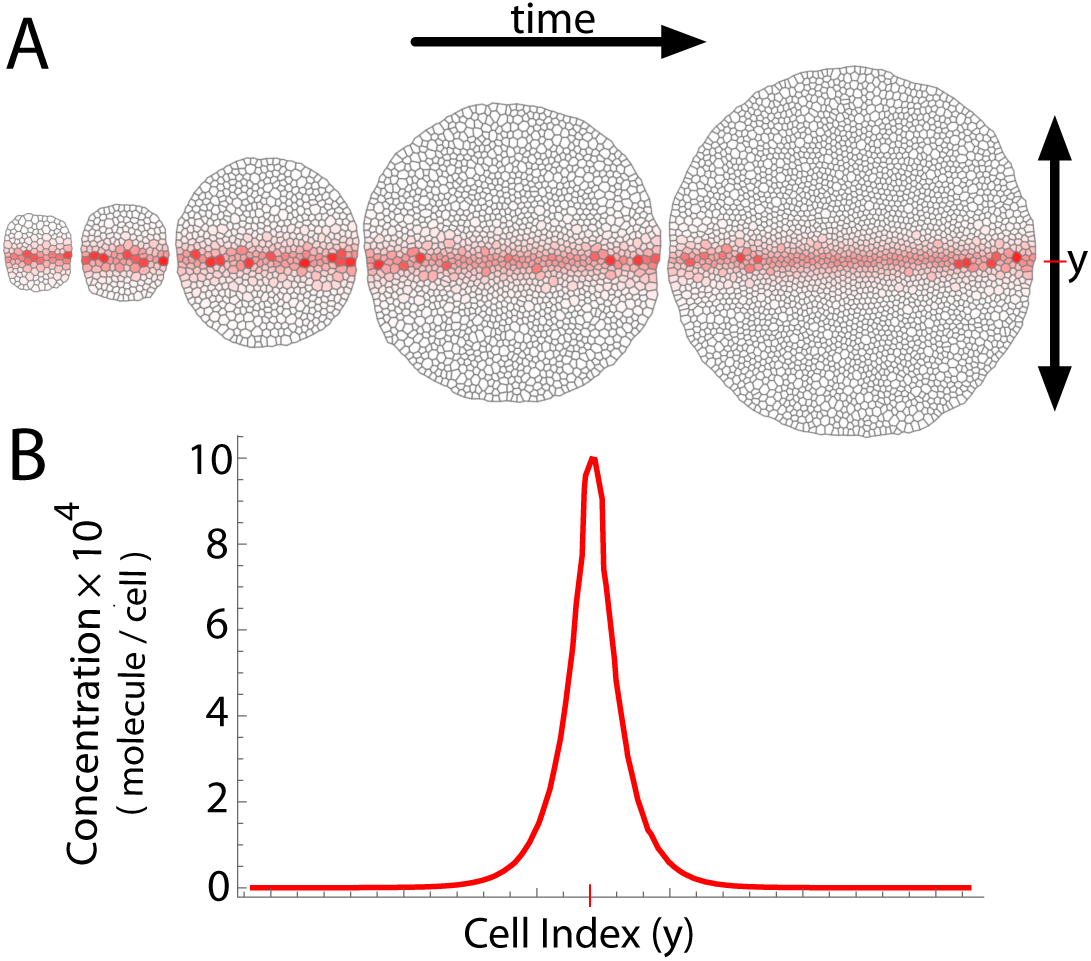
**(Figure S2)** (**A**) Time evolution of a morphogen concentration gradient in the tissue where cells actively grow and divide. The profile reaches a stationary state, **B**.

### Simulation Code: Compilation Instructions

Data and simulation codes for generating each of the figures and movies of the study are provided within the folder `CODE`. The `SRC` folders contain the required C++ source files to generate an executable code when compiled with a standard C++ compiler. In our case the code was compiled in a Linux system using g++. The name of the folders indicates the corresponding figure and/or movie in the main or supplementary text. The executable file is generate through a `Makefile` (i.e., invoking the `make` command within the directory containing the source code).

### Simulation Code: Data Files Output

As a result of a particular simulation different output files are generated. For each simulation, the relevant output files are collected in the `DATA` folder. The output file `dcells.dat` contains the information of all tissue cells for every frame registered in the simulation. The structure of each frame is as follows:

~~~
#cells stageidx
 cellID type #vertexes area #proteinspecies #protein1 #protein2 … #proteinN centerx centery idxneighbor1
idxneighbor2 … idxneighborN vertex1x vertex1y vertex2x vertex2y … vertexNx vertexNy
 cellID type #vertexes area #proteinspecies #protein1 #protein2 … #proteinN centerx centery idxneighbor1
idxneighbor2 … idxneighborN vertex1x vertex1y vertex2x vertex2y … vertexNx vertexNy
 .
 .
 .
 cellID type #vertexes area #proteinspecies #protein1 #protein2 … #proteinN centerx centery idxneighbor1
idxneighbor2 … idxneighborN vertex1x vertex1y vertex2x vertex2y … vertexNx vertexNy
~~~

**Figure 8.**
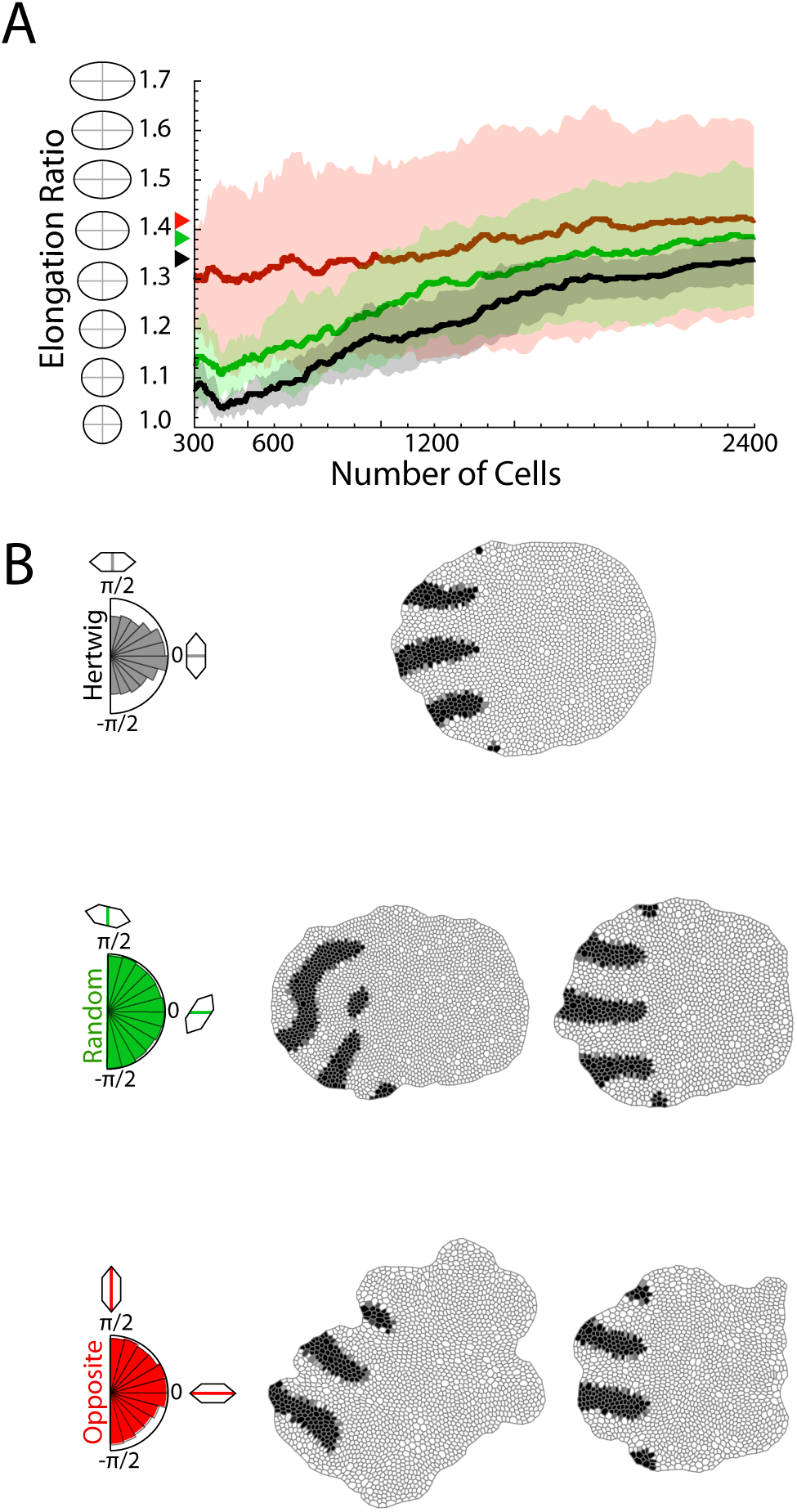
(Figure S3) Effect of cleavage orientation in Turing patterned system. **A**: Tissue elongation as a function of the number of cells for different cleavage dynamics (ten simulations): solid lines stand for the average and the shadings for the standard deviation bands. Hertwig rule leads to the smallest variability indicating a more robust elongation and regular shape. **B**: The polar histograms of cleavage orientations (inset) are a readout of the cellular geometry when the Hertwig rule (black) or its opposite (red) apply but not in the case of random cleavage (green). Final snapshots of representative simulations depending on the cleavage dynamics. Two snapshots for opposite and random cleavage are shown that corresponds to cases where the elongation ratio is smaller/larger than average. Deviations from the Hertwig rule leads to irregular tissue shapes.

That is, each frame starts with a single line that indicated the number of cells in that frame and the stage index. The former, in turn, indicates the number of lines for each frame. Each of those lines contains information of a cell in the following order: the ID of the cell (0, 1, 2…), the type of the cell (1, 2, 3, …), the number of vertexes, the cell area, the number of protein species, their values (number of proteins), the coordinates of the cell center, the indexes of its neighbors, and the coordinates of the cell vertexes (clockwise orientation). For information about the number of protein species and their order for each simulation, see the `protein_order.dat` file that is generated when running the code.

The ID of the cell (`cellID`) is a unique string that is maintained up to a division event: following a division each daughter cell receives a new ID that is the result of joining (by hyphens) the ID of the mother plus and an additional number. Thus, the cell ID allows to reconstruct its lineage: e.g. the cell `405-345-33-67` is a third generation cell (number of hyphens) originating from cell 405 of the initial tissue (its grand-grandmother) and its mother and grandmother are cells `405-345-33` and `405-345` respectively. We point out that the index of a neighbor (e.g. `idxneighbor1`) does not correspond to a *real* ID of the cell (a string such as `405-345-33-67`) but to a number that is the internal ID of the cell in the code. The latter corresponds to the ordinal index of the line within the frame minus 1 (i.e. cell identities counting starts with 0). For example an index of the neighbor `564` corresponds to the cell that is the line `565` in the listing of cells in that frame. The code generates also the output file `divisions.dat` that accounts for the information about cell divisions. Each line of the file reads:

~~~
 stageidx idxduration idxintermediate idxmothercell IDdaughter1 IDdaughter2 type divisionangle
divisionanglehertwig divisionangleplanned centerx centery area #cells #vertexes vertex1x vertex1y vertex2x
vertex2y … vertexNx vertexNy
~~~

Each line indicates the stage index and those of an external loop (`duration`) and an internal loop (`intermediate`) that account for the time evolution of the simulation: each `duration` step contains a given number of `intermediate` steps such that the total (dimensionless) time of a simulation reads (`duration`× `intermediate`×Δ*t* (where Δ*t* is the time step used in the Euler algorithm). Other information provided is the *internal* ID of the mother cell, the *real* IDs of daughter cells, their type, the *actual* division angle, `divisionangle`, the division angle assuming no deviation from the Hertwig rule, `divisionanglehertwig`, the planned division angle, `divisionangleplanned`, the coordinates of the cell center (mother cell), the *actual* cell area at division time (mother cell), the updated number of cells in the tissue, the number of vertexes (mother cell), and the coordinates of those.

Finally, the code generates a log file, `tifosi.log`, where the processes that change the topology of the tissue (e.g., a T1 transition) are stored. The file keeps the time of the event by indicating the `stageidx`, the `idxduration`, and `idxintermediate` and additional information of the process. In the case of a T1 transition, it is provided the internal ID of the edge that disappears, its coordinates (i.e., those of the vertexes that define the edge), the *real* IDs of the cells that shared the edge prior to the T1 transition, the coordinates of the new edges after the transition, and the real IDs of the cells that become connected after the transition takes place. This is an example of how a T1 transition is captured in the log file:

~~~
1 0 3733: t1 process on edge 738 (21.219, 12.1942)-(21.2691, 12.2308) that divides cells 245 and 225.
Coordinates after transtition: (21.2623, 12.1874)-(21.2257, 12.2375) and divides cells 224 and 246.
~~~

In the case of a T2 transition (disappearing triangular cell), besides registering the timing of the event, the log keeps a record of the *real* ID of the triangular cell, the internal ID of the edge that disappears the first (and its coordinates prior to the transition), and also the real IDs of the cell that shared that edge. For example,

~~~
1 2 7452: t2 process on cell 224 triggering edge 736 (19.3435, 14.0312)-(19.2848,14. that divides cells 225 and 224.
~~~

In the case of a T3 transition (two neighboring triangular cells that simultaneously disappear) the information is similar to that of a T2 transition and the log file reads:

~~~
1 23 45632: t3 process on cell 45-345-12 and cell 34 triggering edge 7 (1.3367, 20.4561)-(19.3490, 0.6511)
that divides cells 45-345-12 and 73-465.
~~~

Finally, in the case of a division, the timing and real IDs of the daughter cells are provided, for example,

~~~
1 74 8724: cell division. Daughter cells: 252-401 and 252-400
~~~

### SI Movies

**Movie S1:** The DAH mechanism cannot generate axis extension.

**Movie S2:** Time evolution of a morphogen concentration gradient in the tissue where cells are actively growing and dividing.

**Movie S3:** Simulation of the auto-catalytic intercalation mechanism in a tissue patterned by a morphogen gradient.

**Movie S4:** Simulation of tissue growth if adhesion is not modulated by the morphogen signal.

**Movie S5:** Simulation of tissue growth in the presence of a spatial modulation of the cellular proliferation rates and the auto-catalytic intercalation mechanism in a tissue patterned by stripe alignment of Turing system (Turing patterned tissue).

**Movie S6:** Simulation of tissue growth with spatial modulation of the cellular proliferation rates (Turing patterned tissue).

**Movie S7:** Simulation of tissue growth: control simulations (Turing patterned tissue) without auto-catalytic intercalations or modulation of cellular proliferation.

**Movie S8:** Simulation of tissue growth with auto-catalytic intercalations but without modulation of cellular proliferation (Turing patterned tissue).

**Movie S9:** Cells producing the morphogen responsible of stripe alignment and the modulation of cellular proliferation (Turing patterned tissue).

**Movie S10:** Morphogen concentration profile responsible of stripe alignment and the modulation of cellular proliferation (Turing patterned tissue).

## ACKNOWLEDGMENTS

We thank Dr. Luis M. Escudero (Seville University) for comments on the manuscript. This work was supported by Lehigh University through a Faculty Innovation Grant (FIG-2019-JB).

